# *Ruegeria persistens* sp. nov., a persistent symbiotic bacterium isolated from coral

**DOI:** 10.64898/2026.09.01.748717

**Authors:** Mei Xie, Congjuan Xu, Christian R. Voolstra, Haiwei Luo

**Author notes:** **Corresponding author:** Haiwei Luo, School of Life Sciences, The Chinese University of Hong Kong, Shatin, Hong Kong SAR.

## Abstract

A Gram-stain-negative, aerobic, moderately halophilic, non-motile, rod-shaped bacterium, designated MC10-B4^T^, was isolated from the coral *Oulastrea crispata* collected from Hong Kong waters. The strain showed the ability to persist in a coral holobiont during an eight-month monitoring period in a field trial, indicating a long-term association with the coral host. Growth occurred at 10-37 °C (optimum, 28-30 °C), at NaCl concentrations up to 4.0% (w/v; optimum, 0.5-2.0%), and at pH 7.0-10.0 (optimum, pH 7.0-8.0) under standard laboratory conditions. Colonies on Marine Agar 2216 were round, smooth, convex, white, opaque, and had entire margins. Phylogenetic analysis based on 16S rRNA gene sequences indicated that strain MC10-B4^T^ belongs to the genus *Ruegeria* and shares the highest sequence similarity with *Ruegeria conchae* TW15^T^ (99.64%), followed by *Ruegeria profundi* ZGT108^T^ (98.92%), *Ruegeria lacuscaerulensis* ITI-1157^T^ (98.46%), *Ruegeria denitrificans* CECT 5091^T^ (98.31%), *Ruegeria arenilitoris* G-M8^T^ (97.69%), *Ruegeria atlantica* NBRC 15792^T^ (97.55%), *Ruegeria halocynthiae* MA1-6 (97.11%), *Ruegeria atlantica* IAM 14463^T^ (96.98%), *Ruegeria meonggei* MA-E2-3 (96.83%), and *Ruegeria pomeroyi* DSS-3^T^ (96.65%). Genome sequencing revealed a genome size of 4.55 Mbp and a DNA G+C content of 57.5 mol%. The average nucleotide identity and digital DNA-DNA hybridization values between strain MC10-B4^T^ and *Ruegeria conchae* TW15^T^ were 77.6% and 18.2-22.8%, respectively. These values are below the recommended thresholds for species delineation (95% and 70%). Strain MC10-B4^T^ shares the same predominant fatty acid, summed feature 8 (consisting of C_18:1_ *ω*7*c* and/or C_18:1_ *ω*6*c*), with other *Ruegeria* strains; however, it also contains C_19:0_ cyclo *ω*8*c*, which was not detected in *Ruegeria conchae* TW15^T^ and *Ruegeria pomeroyi* DSS-3^T^. The type strain is MC10-B4^T^ (= GDMCC 1.6889^T^ = KCTC 18785^T^).

## Introduction

Bacteria of the genus *Ruegeria* (family *Rhoseobacteraceae*, class *Alphaproteobacteria*) are frequently detected in coral holobionts across diverse geographic regions [1-4]. *Ruegeria* species utilize a range of coral-derived osmolytes and host-associated substrates, which is consistent with adaptation to the coral niche [5, 6]. Although many host-associated bacteria are considered transient opportunists, some strains form persistent associations with their hosts and may contribute to host performance [7, 8]. The taxonomic identity and functional capacity of such persistent coral-associated *Ruegeria* strains have remained poorly characterized.

A recent evolution-guided screening of coral-associated bacterial isolates identified a *Ruegeria* population designated MC10 that exhibits genomic hallmarks of early-stage host dependency, including insertion sequence proliferation and pseudogenization of core metabolic pathways [9]. A strain from this population, MC10-B4, demonstrated sustained colonization in the coral *Acropora pruinosa* over an eight-month field trial and was associated with enhanced host thermal resilience during a natural bleaching event [10]. These findings raise the question of whether strain MC10-B4 represents a previously undescribed species within the genus *Ruegeria*.

Strain MC10-B4^T^ was isolated from the coral *Oulastrea crispata* collected from Hong Kong waters. Phylogenetic analysis based on 16S rRNA gene sequences indicated that strain MC10-B4^T^ belongs to the genus *Ruegeria* and shares the highest sequence similarity with *Ruegeria conchae* TW15^T^ (99.64□%), followed by *Ruegeria profundi* ZGT108^T^ (98.92□%) and *Ruegeria denitrificans* CECT 5091^T^ (98.31□%). The average nucleotide identity (ANI, 77.6□%) and digital DNA-DNA hybridization (dDDH, 18.2–22.8□%) values between strain MC10-B4^T^ and *R. conchae* TW15^T^ are below the recommended thresholds for species circumscription (95□% and 70□%, respectively) [11], which indicates that strain MC10-B4^T^ represents a distinct species from known members of the genus *Ruegeria*.

Phenotypic and chemotaxonomic characterization revealed that strain MC10-B4^T^ is Gram-stain-negative, aerobic, moderately halophilic, non-motile and rod-shaped. The predominant fatty acid is summed feature 8 (C_18:1_ *ω*7*c* and/or C_18:1_ *ω*6*c*), which is shared with other *Ruegeria* strains, but strain MC10-B4^T^ also contains C_19:0_ cyclo *ω*8*c*, a feature not detected in *R. conchae* TW15^T^ or *Ruegeria pomeroyi* DSS-3^T^. The strain lacks detectable catalase activity and is oxidase-negative, does not hydrolyse Tween 80 or casein, and lacks motility. The DNA G+C content is 57.5 mol%. Based on the evidence presented in this study, strain MC10-B4^T^ is assigned to the genus *Ruegeria* and is proposed as the type strain of a new species, *Ruegeria persistens* sp. nov.

### Isolation and cultivation

Strain MC10-B4^T^ was isolated from the coral *Oulastrea crispata* collected from Bluff Island in Hong Kong waters at a depth of 2-3.5 meters. Coral fragments were collected using a hammer and chisel and shipped to the laboratory at the Chinese University of Hong Kong for processing in the same day. Coral fragments (∼5 cm^2^) were washed three times with autoclaved 0.22 μm-filtered seawater (AFSW) to remove loosely attached microbes on coral surfaces. Mucus was collected using a 100 μL pipette, serially diluted (10-fold), and spread on marine basal medium (MBM) agar plates. The medium recipe was modified from a previous protocol [12] by replacing the nitrogen source with glycine betaine and N-acetylglucosamine to achieve a higher *Ruegeria* isolation efficiency. Plates were incubated at 28 °C for two weeks, and single colonies were picked, purified three times on Marine Agar 2216 (BD Difco, USA) and stored in 25% glycerol at -80 °C.

### 16S rRNA gene sequence and phylogeny

The taxonomic affiliation of strain MC10-B4^T^ was initially assessed via full-length 16S rRNA gene sequence analysis. BLASTn searches [13] against the EzBioCloud database [14] revealed that strain MC10-B4^T^ is a member of the genus *Ruegeria*. The strain exhibited the highest 16S rRNA gene sequence similarity to *R. conchae* TW15^T^ (99.64%), followed by *R. profundi* ZGT108^T^ (98.92%) and *R. denitrificans* CECT 5091^T^ (98.31%).

To confirm the taxonomic placement, a maximum-likelihood phylogenetic tree was reconstructed. The complete 16S rRNA gene sequence of strain MC10-B4 ^T^ was obtained from the complete genome. Reference sequences included: (i) all 23 validly published *Ruegeria* species based on LPSN (https://lpsn.dsmz.de/genus/ruegeria) [15] plus *Ruegeria* sp. 6PALISEP08 and *Ruegeria* sp. AD91A. *R. gelatinovorans* was excluded as its 16S rRNA gene sequence was not available. (ii) 15 additional type strains representing related genera within the Rhodobacteraceae family (*Sulfitobacter, Aliishimia, Epibacterium, Pseudooceanicola, Albibacillus, Roseovarius, Thalassovita, Leisingera, Phaeobacter, Thiosulfatihalobacter, Shimia*). All reference sequences were retrieved from the EzBioCloud database [14].

Multiple sequence alignment was performed using MAFFT v7.471 with the --auto parameter [16], and the resulting alignments were trimmed using trimAl v1.4 with the “automated1” mode [17]. A maximum-likelihood (ML) phylogenetic tree was reconstructed using IQ-TREE v1.6.12 with the best-fit nucleotide substitution model automatically selected by ModelFinder [18]. Branch support was assessed using 1,000 ultrafast bootstrap replicates. The tree was edited and displayed with iTOL v5 [19].

In the phylogenetic tree (**Fig. 1**), MC10-B4^T^ belonged to the genus *Ruegeria* and had the highest similarities to *R. conchae* TW15^T^. Crucially, strain MC10-B4^T^ was further delineated from its closest relative, *R. conchae* TW15^T^, by ANI and dDDH values of 77.6% and 18.2–22.8%, respectively, both falling well below the recommended thresholds for species circumscription [11].

**Fig. 1.**
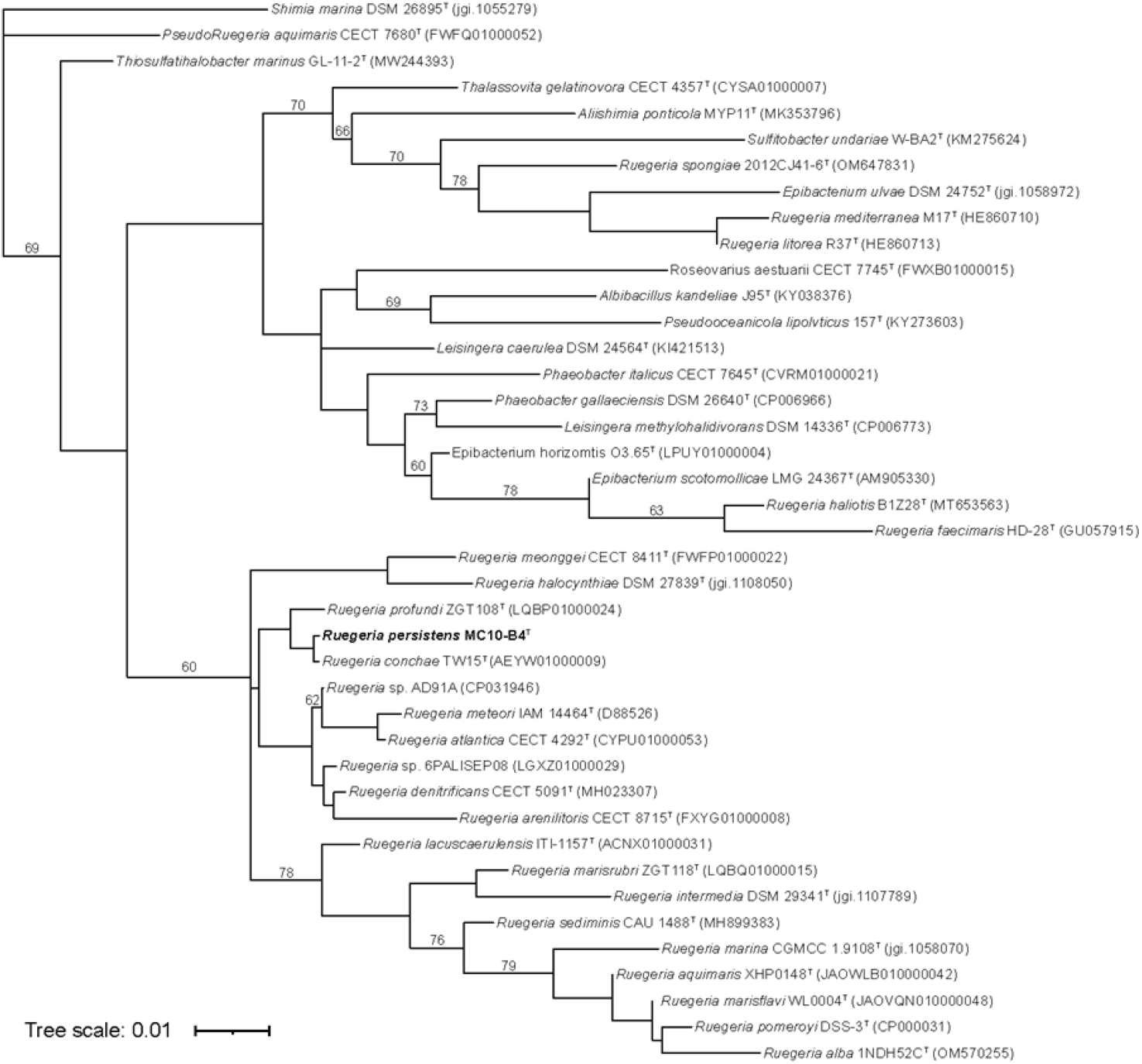
Maximum-likelihood phylogenetic tree based on 16S rRNA gene sequences showing the relationships between strain MC10-B4^T^ and the type strains of recognized *Ruegeria* species. The tree was reconstructed using IQ-TREE with the best-fit nucleotide substitution model automatically selected by ModelFinder. Bootstrap values (1,000 ultrafast replicates) are shown at the nodes. Bootstrap values ≥60% are shown at branch points. Sequences from 15 additional type strains representing related genera within the family *Rhodobacteraceae* were included as outgroups. Bar, 0.01 substitutions per nucleotide position.

### Morphology, physiological, and biochemical characterization

A Gram stain kit (Huankai Microbial, China) was used to determine the Gram reaction. Cell morphology was observed using a transmission electron microscope (Hitachi H-7650, Japan). Swimming, swarming, and twitching motilities were tested on Marine 2216 agar plates at 28 °C, with swimming motility tested on plates containing 0.18% agar and incubated for 6 days; swarming motility tested on plates containing 0.75% agar and incubated for 10 days; twitching motility tested on plates containing 1% agar and stab-inoculated, then incubated for 10 days. To determine the optimal growth conditions of the strain MC10-B4^T^, growth was investigated in Marine Broth (MB) 2216 medium (BD Difco, USA) at various temperatures (4, 10, 15, 20, 25, 28, 30, 37, 40, 42, 45 and 50 °C), NaCl concentrations (0, 0.5, 1.0, 2.0, 3.0, 4.0, 5.0, 6.0, 7.0, and 10.0%, w/v) and pH levels (pH 4.0-11.0, at 1.0 pH unit intervals). Catalase activity was examined by observing bubble production using 5.0% hydrogen peroxide. Oxidase activity was determined by the commercial Oxidase reagent kit (Huankai Microbial, China). Enzymatic activities and biochemical characteristics were determined using the API ZYM kit and the API 20NE system. The API ZYM kit detects the hydrolysis of chromogenic or fluorogenic substrates. The API 20NE system assesses the assimilation of 12 carbon sources and includes the following physiological tests: nitrate reduction, indole production, urease, esculin hydrolysis, gelatinase, β-galactosidase, glucose fermentation, and arginine dihydrolase.

The physiological and biochemical characteristics of strain MC10-B4^T^ and the related *Ruegeria* type strains are presented in Table 1. Strain MC10-B4^T^ was Gram-stain-negative, catalase-weakly positive, and oxidase-negative. It did not hydrolyze Tween 80 or casein. The cells of strain MC10-B4^T^ are approximately 0.6-0.7 μm wide and 1.1-1.7 μm long. Transmission electron microscopy revealed rod-shaped cells containing prominent electron-lucent storage granules, consistent with polyhydroxyalkanoates (PHAs) (**Fig. 2a**). Light microscopy following Gram staining confirmed the Gram-negative reaction and rod morphology (**Fig. 2b**). Strain MC10-B4^T^ exhibited weak swimming motility (zone diameter: 1.9 ± 0.1 cm) but no swarming or twitching motility. Growth occurred at 10-37 °C (optimum, 28-30 °C), at NaCl concentrations up to 4.0% (w/v; optimum, 0.5-2.0%) and at pH 7.0-10.0 (optimum, pH 7.0-8.0) under the standard laboratory conditions.

**Table 1.** Physiological and biochemical characteristics distinguishing strain MC10-B4^T^ from closely related species of the genus *Ruegeria*. Strains: 1, MC10-B4^T^; 2, *R. pomeroyi* DSS-3^T^; 3, *R. conchae* TW15^T^; 4, *R. profundi* ZGT108^T^; 5, *R. halocynthiae* KCTC 23463^T^; 6, *R. arenilitoris* G-M8^T^; +, positive; w, weakly positive; -, negative. Data taken from: a, Gonzalez et al., 2024 [20]; b, Lee et al., 2012 [21]; c, Zhang et al., 2017 [22]; d, Kim et al., 2012 [23]; e, Park et al., 2012 [24].

| Characteristic | 1 | 2 | 3 | 4<sup>c</sup> | 5<sup>d</sup> | 6<sup>e</sup> |
| --- | --- | --- | --- | --- | --- | --- |
| Isolation source | Coral | Seawater<sup>a</sup> | Ark clam<sup>b</sup> | Brine-seawater interface | Sea squirt | seashore sand |
| Temperature range (optimum) (°C) | 10-37 (28-30) | 10-40 (30)<sup>a</sup> | 10-37 (25-30)<sup>b</sup> | 15-42 (33-35) | 10-37 (30) | 4-45 (30-37) |
| pH range (optimum) | 7.0-10.0 (7.0-8.0) | ND<sup>a</sup> | 7.0-10.0 (8.0)<sup>b</sup> | 5.5-9.0 (7.5-8.0) | 5.5-8.0 (7.0-8.0) | 5.5-8.0 (7.0-8.0) |
| NaCl (optimum) (% w/v) | 0.5-4.0 (0.5-2.0) | 1.5-7.0 (0.6-2.3)<sup>a</sup> | 1.0-5.0 (2.0)<sup>b</sup> | 2.9-11.1 (5.8-8.8) | 2.0-6.0 (2.0-3.0) | 0.5-6.0 (2.0) |
| Motility | w | +<sup>a</sup> | -<sup>b</sup> | - | - | + |
| Major fatty acids (>10%) | Summed features 8; C<sub>18:1</sub> ω7c 11-Methyl | C<sub>18:1</sub> ω7c, C<sub>16:0</sub><sup>a</sup> | Summed features 8; C<sub>18:1</sub> ω7c 11-Methyl<sup>b</sup> | Summed features 8; C<sub>16:0</sub> 2OH | C<sub>18:1</sub> ω7c, C<sub>18:1</sub> ω7c 11-Methyl | C<sub>18:1</sub> ω7c, C<sub>18:1</sub> ω7c 11-Methyl |
| **API ZYM test:** |  |  |  |  |  |  |
| Alkaline phosphatase | + | + | + | + | + | + |
| Esterase (C4) | + | + | + | + | w | w |
| Esterase lipase (C8) | + | + | + | + | + | - |
| Lipase (C14) | w | w | w | - | - | - |
| Leucine arylamidase | + | + | + | + | + | w |
| Valine arylamidase | w | + | w | - | - | - |
| Cystine arylamidase | - | - | w | - | - | - |
| Trypsin | - | - | - | - | - | - |
| Chymotrypsin | - | - | - | - | - | - |
| Acid phosphatase | + | + | + | w | + | - |
| Naphthol-AS-BI-phosphohydrolase | + | + | + | - | - | - |
| α-galactosidase | - | - | - | - | - | - |
| β-galactosidase | - | - | w | - | - | - |
| β-glucuronidase | - | - | - | - | - | - |
| α-Glucosidase | - | - | w | w | + | - |
| β-glucosidase | - | - | - | - | - | - |
| N-Acetyl-β-glucosaminidase | - | - | - | - | - | - |
| α-Mannosidase | - | - | w | - | - | - |
| α-Fucosidase | - | - | - | - | - | - |
| **API 20 NE tests:** |  |  |  |  |  |  |
| Reduction of nitrates to nitrites | - | - | - | - | + | + |
| Esculin | w | w | w | + | - | - |
| Gelatin | - | w | - | - | - | - |
| D-Mannose | - | - | - | w | + | + |
| D-Mannitol | - | - | - | + | - | - |
| Malate | - | - | - | - | + | + |
| Citrate | - | - | - | - | w | + |
| DNA G+C content (mol%) | 57.5 | 68.0 <sup>a</sup> | 55.7 <sup>b</sup> | 56.7 | 58.6 | 64.6 |

**Fig. 2.**
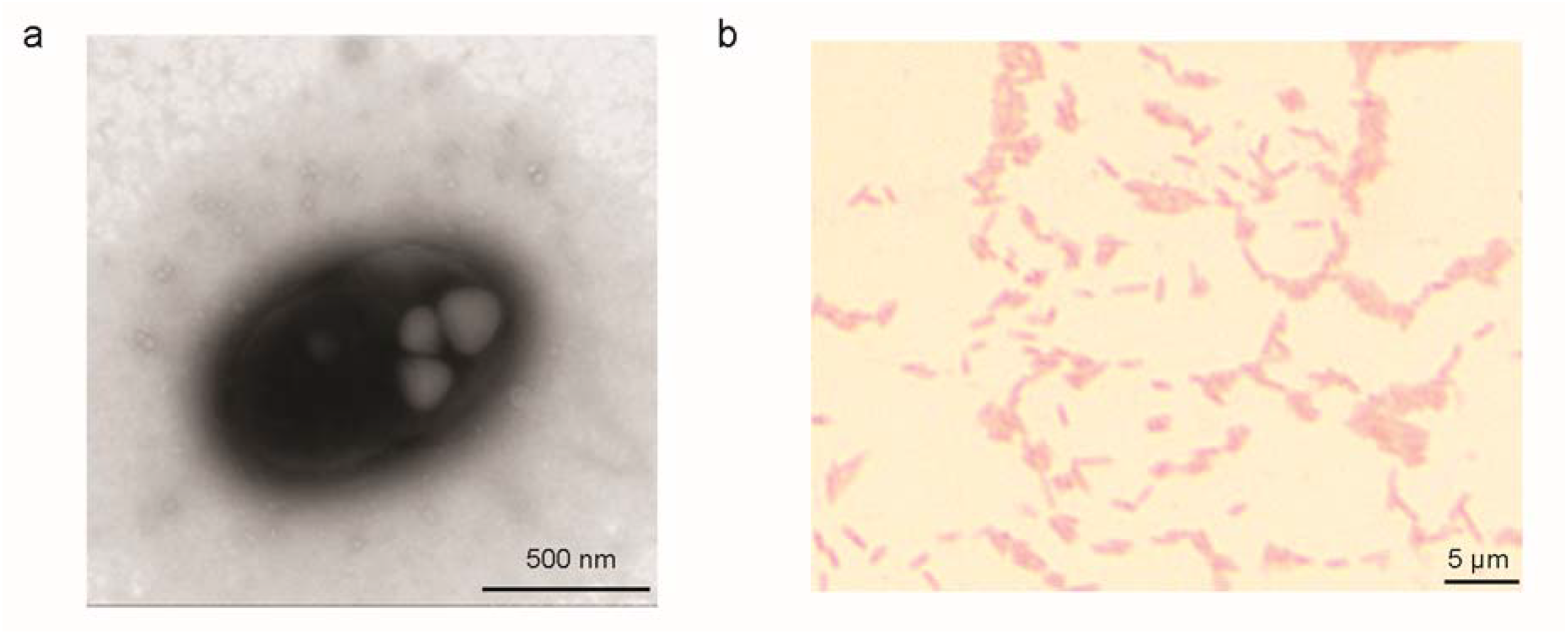
Morphology of strain MC10-B4^T^. (a) Transmission electron micrograph showing cell ultrastructure. Bar, 500 nm. (b) Light micrograph following Gram staining, indicating Gram-negative reaction (pink) and rod-shaped cell morphology. Bar, 20 μm.

In API ZYM tests, strain MC10-B4^T^ showed positive activities for alkaline phosphatase, esterase (C4), lipase esterase (C8), leucine arylamidase, acid phosphatase, and naphthol-AS-BI-phosphohydrolase, and weak activity for lipase (C14) and valine arylamidase. All other enzymes tested were negative. In API 20NE tests, strain MC10-B4^T^ was negative for all tests except weak positive esculin hydrolysis, with no carbon source assimilation detected. Strain MC10-B4^T^ was clearly differentiated from *R. conchae* TW15^T^ (lacking cystine arylamidase, β-galactosidase, α-glucosidase, and α-mannosidase activities) and *R. pomeroyi* DSS-3^T^ (weaker valine arylamidase and negative gelatin hydrolysis).

### Chemotaxonomic characterization

The cellular fatty acid profile of strain MC10-B4^T^ was analyzed using the Sherlock Microbial Identification System (MIDI, Inc., Newark, DE, USA) RTSBA6 method. Strain MC10-B4^T^ was grown on Marine 2216 agar at 28 °C for 48 h. The fatty acid profiles were analyzed by gas chromatography, and the resulting peaks were identified using the MIDI Sherlock software (version 6.1). The analysis met the system’s quality control standards, with an ECL deviation of 0.003 (acceptable limit <0.020) and a reference ECL shift of 0.002 (acceptable limit <0.010), based on five reference peaks. The total response was 721,968, of which 718,164 (99.47%) were successfully identified.

The predominant fatty acids were summed feature 8 (C_18:1_ *ω*7c / C_18:1_ *ω*6c, 40.60%) and C_18:1_ *ω*7c 11-methyl (19.18%). Substantial amounts of hydroxylated fatty acids were also detected: 16:0 2OH (12.83%), 12:0 3OH (6.75%), and 19:0 cyclo *ω*8c (7.00%). Saturated straight-chain fatty acids included 10:0 (3.51%), 12:0 (4.31%), 16:0 (2.15%), and 18:0 (1.04%). Minor components comprised 18:1 2OH (1.52%), 17:0 (0.09%), 16:1 2OH (0.27%), 17:0 2OH (0.22%), 18:0 2OH (0.12%), 20:1 *ω*7c (0.16%), 20:2 *ω*6,9c (0.06%), 15:0 2OH (0.08%), and summed feature 3 (0.11%). The presence of 19:0 cyclo *ω*8c (7.00%) is a distinguishing feature of strain MC10-B4^T^, which was not detected in either *R. conchae* TW15^T^ or *R. pomeroyi* DSS-3^T^. Cyclopropane fatty acids are often associated with stress responses or stationary phase growth in bacteria [25, 26].

Polar lipids were extracted from freeze-dried cells of strains MC10-B4^T^ and *R. pomeroyi* DSS-3^T^ and separated by two-dimensional thin-layer chromatography (TLC) on silica gel plates. The first dimension used chloroform: methanol: distilled water (65:25:4, v/v), and the second dimension used chloroform: glacial acetic acid: methanol: distilled water (80:18:12:5, v/v). Lipids were visualized using phosphomolybdate (total lipids), ninhydrin (aminolipids), reagent D (phospholipids), molybdenum blue (phosphate-containing lipids), and 1-naphthol (glycolipids). Both strains shared the major phospholipids phosphatidylglycerol (PG), phosphatidylethanolamine (PE), and phosphatidylcholine (PC). Compared to *R. pomeroyi* DSS-3^T^, which contained five unidentified phospholipids (PL1– 5), two aminolipids (AL1–2), and two aminophospholipids (APL1–2), strain MC10-B4^T^ exhibited a simpler profile with only two unidentified phospholipids (PL1–2), one aminolipid (AL), and no detectable aminophospholipids. Both strains contained two unidentified lipids (L1–2).

Respiratory quinones were extracted from freeze-dried cells of strains MC10-B4^T^ and *R. pomeroyi* DSS-3^T^ using chloroform/methanol (2:1, v/v). Purified quinones were analyzed by reversed-phase HPLC with methanol/isopropyl ether (2:1, v/v) as the mobile phase and detected at 254 nm. Identification was based on comparison of retention times with authentic standards (Q-10 and MK-7). HPLC analysis revealed that strain MC10-B4^T^ contains Q-10 as the sole respiratory quinone (retention time: 20.872 min; peak area: 2,887,527; purity: 100%). In contrast, *R. pomeroyi* DSS-3^T^ contains both Q-10 (retention time: 20.883 min; peak area: 2,447,243; purity: 100%) and MK-7 (retention time: 15.173 min; peak area: 2,551,958; purity: 100%).

### Genomic and functional characterization

The complete genome sequence of strain MC10-B4^T^ was determined using a hybrid assembly approach combining short-read sequencing on the DNBSEQ PE150 platform (BGI, China) and long-read sequencing on the Oxford Nanopore Technologies MinION platform with an R9.4.1 flow cell. The genome assembly was circularized into a single chromosome of 4.55□Mbp. The DNA G+C content is 57.5□mol%. The raw reads from short-read and long-read sequencing, as well as the assembled genome sequence of strain MC10-B4^T^, have been deposited in GenBank under accession numbers SRR33968895 (short reads, https://trace.ncbi.nlm.nih.gov/Traces/?run=SRR33968895), SRR33961571 (long reads, https://trace.ncbi.nlm.nih.gov/Traces/?run=SRR33961571), and GCA_056535805.1 (assembly, https://www.ncbi.nlm.nih.gov/datasets/genome/GCF_056535805.1/).

Comparative genomic analysis against the closest phylogenetic relative, *Ruegeria conchae* TW15^T^, revealed average nucleotide identity (ANI) of 77.6□% and digital DNA-DNA hybridization (dDDH) values ranging from 18.2□% to 22.8□%, both well below the species circumvention thresholds of 95□% and 70□%, respectively [11]. These values confirm that strain MC10-B4^T^ represents a distinct species.

The genome of strain MC10-B4^T^ exhibits several features consistent with an emerging host-associated lifestyle. Insertion sequences have proliferated, with multiple copies across the chromosome, and pseudogenes have accumulated, including disruptions in core metabolic pathways such as the tricarboxylic acid cycle [10]. Genes encoding siderophore biosynthesis (*entABCDEFS*) and exopolysaccharide production (*exo* operon) are present, and these genomic predictions were supported by phenotypic assays showing iron chelation and biofilm formation [9]. The strain lacks detectable catalase activity, which is consistent with the absence of a functional catalase gene expression under standard laboratory conditions, although catalase genes (*katE, katG*) are present in the genome [9]. For oxidative stress defence, the strain carries a complete cytochrome bd ubiquinol oxidase cluster (*cydABDX*) and a Class Ib ribonucleotide reductase operon (*nrdFHI*), both of which are enriched in the MC10 population compared to other coral-associated *Ruegeria* lineages [9].

### Description of *Ruegeria persistens* sp. nov

*Ruegeria persistens* sp. nov. (L. part. adj. *persistens*, persistent, enduring; referring to the ability of the strain to establish long-term persistence within its host).

Cells are Gram-stain-negative, non-motile rod-shaped, 0.6-0.7 μm wide and 1.1-1.7 μm long. Colonies on Marine Agar 2216 are round, smooth, convex, white, opaque, with entire margins. Temperature range 10-37°C (optimum 28-30°C), NaCl range 0-4.0% (w/v) (optimum 0.5-2.0%), pH range 7.0-10.0 (optimum 7.0-8.0). Catalase weakly positive, oxidase negative. Does not hydrolyse Tween 80 or casein. Positive for alkaline phosphatase, esterase (C4), esterase lipase (C8), leucine arylamidase, acid phosphatase, naphthol-AS-BI-phosphohydrolase (API ZYM); weakly positive for lipase (C14), valine arylamidase, and esculin hydrolysis (API 20NE). No carbon source assimilation (API 20NE). Major fatty acids (>10%) are summed feature 8 (C_18:1_ *ω*7c / C_18:1_ *ω*6c, 40.6%) and C_18:1_ *ω*7c 11-methyl (19.2%). Other fatty acids include C_16:0_ 2OH 2-OH (12.8%), C_19:0_ cyclo *ω*8*c* (7.0%), C_12:0_ 3-OH (6.8%). Polar lipids are phosphatidylethanolamine, phosphatidylglycerol, phosphatidylcholine, two unidentified phospholipids, one unidentified aminolipid, and two unidentified lipids. The predominant respiratory quinone is Q-10. The DNA G+C content of the type strain is 57.5 mol%.

The type strain, MC10-B4^T^ (= GDMCC 1.6889^T^ = KCTC 18785^T^), was isolated from the coral *Oulastrea crispata* collected from Hong Kong waters and has been deposited in the GDMCC (Guangdong Microbial Culture Collection Center) with accession number GDMCC 1.6889 and KCTC (Korean Collection for Type Cultures) with accession number KCTC 18785. The complete genome sequence and 16S rRNA gene sequence of strain MC10-B4^T^ have been deposited at GenBank/EMBL/DDBJ under accession numbers GCA_056535805.1 (https://www.ncbi.nlm.nih.gov/datasets/genome/GCF_056535805.1/) and PZ369073 (https://www.ncbi.nlm.nih.gov/nuccore/PZ369073.1/), respectively.

## Acknowledgements

This study is supported by the Hong Kong Research Grants Council (RGC) General Research Fund (project #: 14114724) and the Coral Research and Development Accelerator Platform (CORDAP) under Award No. CAP-2024-1692.

